# SCRATCH: A programmable, open-hardware, benchtop robot that automatically scratches cultured tissues to investigate cell migration, healing, and tissue sculpting

**DOI:** 10.1101/2024.08.27.609782

**Authors:** Yubin Lin, Alexander Silverman-Dultz, Madeline Bailey, Daniel J. Cohen

## Abstract

**Despite the widespread popularity of the ‘**scratch assay’, where a pipette is dragged through cultured tissue to create an injury gap to study cell migration and healing, the manual nature of the assay carries significant drawbacks. So much of the process depends on individual manual technique, which can complicate quantification, reduce throughput, and limit the versatility and reproducibility of the approach. Here, we present a truly open-source, low-cost, accessible, and robotic scratching platform that addresses all of the core issues. Compatible with nearly all standard cell culture dishes and usable directly in a sterile culture hood, our robot makes highly reproducible scratches in a variety of complex cultured tissues with high throughput. Moreover, we demonstrate how scratching can be programmed to precisely remove areas of tissue to sculpt arbitrary tissue and wound shapes, as well as enable truly complex co-culture experiments. This system significantly improves the usefulness of the conventional scratch assay, and opens up new possibilities in complex tissue engineering and cell biological assays for realistic wound healing and migration research.

## Introduction

The ‘scratch assay’ (Fig. 1A) —dragging a pipette tip or sharp object through a cultured tissue and monitoring the cellular healing response into the resulting gap—is among the most common approaches to study cell migration and healing *in vitro* (1,2), but also perhaps among the least reproducible and scalable due to the manual nature of the process (2–5). While a popular protocol paper on the manual method has nearly 5000 citations at this point (1) and the method is largely free, the traditional scratch assay relies on pressure, tool orientation and brand, speed, and manual stability, and is inherently limited in precision, throughput, and scalability (e.g. it is more difficult in a 96-well plate than in 6-well plate). Moreover, there is a missed opportunity to use ‘scratching’ as a form of subtractive manufacturing to produce much more complex tissue geometries and easily prepare unique systems-level co-cultures. Given the ubiquity and importance of scratch assays, new approaches improving the reproducibility, throughput, and versatility can benefit a broad range of research fields.

**Figure 1.**
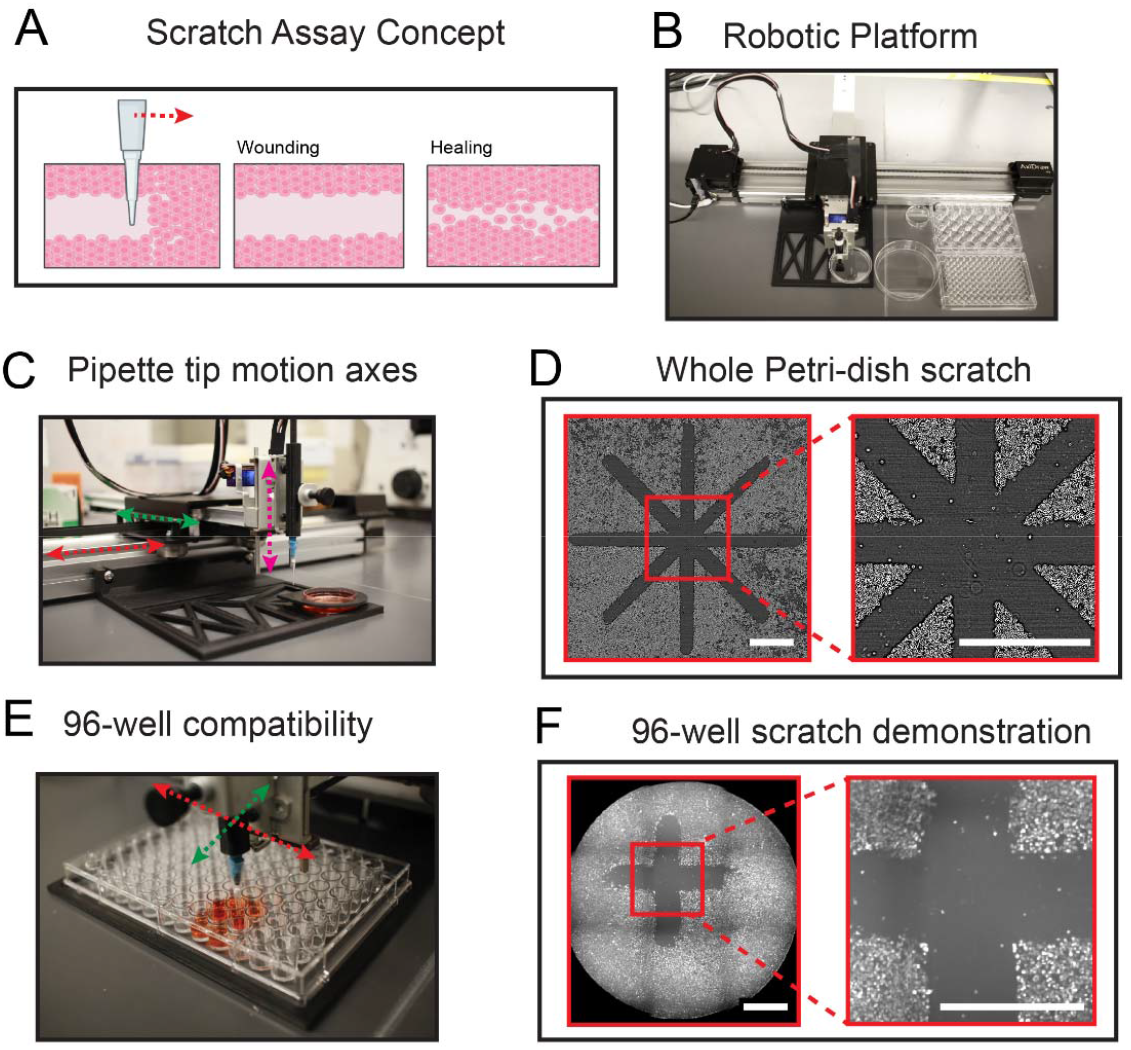
System mechanism and capability. (A) Scratch assay is performed by a pipette tip moving across the cell monolayer, leaving a cell-depleted region. (B) System overview. The lateral movement of the pipette tip is actuated by a stepper motor-driven belt sytem. Red and green arrow represent X and Y direction respectively. The vertical movement of the tip is actuted by a servo, indicated with magenta arrow. The 35mm dish is placed on a custom designed fixture. (C) A close-up photo of the device operating on a 60mm dish. (D) Phase-contrast image of dish-scale scratch pattern demonstration, scale bar: 2mm. (E) Cytoplasmic staining of scratch pattern in a 96-well dish, scale bar: 1mm. (F) Close-up photo of device operating on a 96-well plate. Arbitrary pattern and well location can be selected.

While alternative solutions to generate gaps in tissues are well-represented in the literature, none of them address all of the challenges (2). One popular approach is the ‘barrier removal assay’ where cells are seeded on either side of a rubber stencil and then the stencil is removed to generated a ‘gap’ (6–9). While versatile, the approach requires precision pipetting (10), and simply does not scale to small culture vessels. Commercial rubber inserts are available, but are limited in geometry and configuration as well as being costly consumables. Further, there is a concern that barrier removal does may not properly damage the surrounding tissue consistent with actual injury (11). Similarly, DIY parallel scratchers based on machined or molded tips have been effectively used in multi-well plate studies (3,12,13), but the approach relies on sophisticated machine shop CNC capabilities, still requires user applied pressure and speed, can only make straight lines, and is intrinsically limited to a single specific substrate (e.g. 96-well only). While commercial scratch systems exist (14,15), they are also limited to only a few well-plates options (e.g. 24/96-well) and straight lines, and the cost is prohibitively high, relatively speaking (∼10k-20k USD at the time of writing). Finally, numerous non-mechanical strategies have been developed that rely on electrical, chemical, and optical patterning allowing improved precision (down to the micron scale), but carry their own limitations to cost, throughput, and versatility (2). Hence, there is an exciting opportunity to redevelop the common, mechanical form of the scratch assay both around flexible, programmable, open-hardware that can be adopted by any laboratory.

All of the key variables and challenges discussed here are the things that a robot excels at—precision, reproducibility, throughput/repetition, and programmability. Inspired by these advantages, we modified a low-cost robotic platform originally intended for art generation. We call this device SCRATCH—Scalable Cellular Resection Apparatus To Characterize Healing. SCRATCH allows: (1) complete programmability to produce almost any pattern; (2) the use of any scratching tip (e.g. pipette tips, needles, wires, etc.); (3) compatibility with nearly all standard culture vessels (3.5 cm dishes to 96-well plates); (4) direct use in a sterile culture hood; and (5) a low net cost of <500USD at the time of writing (16). The remainder of this report summarizes how SCRATCH works and demonstrates its capabilities.

## Results

### SCRATCH device working principles and system architecture

SCRATCH is a fully automated scratch assay system, and its key advantages stem from computer-control of a robotic gantry (Fig.1B). The core of the SCRATCH device is a writing/drawing robot that provides programmable lateral (XY) and vertical (Z) movement of the scratching apparatus (Fig. 1C, color-coded arrows). While SCRATCH can be built using off-the-shelf components from the 3D printing community, for simplicity here we modified a hobby ‘art-bot’ (AxiDraw V3, but many others exist) originally intended to hold pens and markers as this saves considerable time for a minimal cost (∼$500). This chassis consists of an XY stepper motor-belt system to position the pipette-tip tool over a tissue culture region, and a servo motor to precisely and gently bring the tool into contact with the tissue in preparation for scratching. Instead of a pen or marker, we 3D-printed a customized pipette tip holder for 10 μL pipette tips (this can be tuned for any pipette tip style) (Fig. 1C). To ensure stability of the tip during scratching, we applied a thin layer of reusable adhesive putty (e.g. FunTak) between the tip and the holder. This tip carrier can then be attached to the XYZ gantry as if it were a pen (see Data Availability for CAD file). At this point, SCRATCH is ready for use (see Video S1 for its operation).

A key design goal was to make SCRATCH as user-friendly and reproducible as possible to enable rapid adoption in cell biology labs, so a key feature of our design is our modular sample-holder directly attached to the frame of SCRATCH that allows most standard culture vessels—from 3.5 cm Petri dishes to 96-well plates (Figs. 1D-F)—to be precisely and reproducibly positioned relative to the pipette tool (see CAD file access instructions in Data Availability; see Fig. S1). This sample holder also incorporates an alignment ring to calibrate the tip position at the beginning of the scratch (see Methods). The use of this fixture allows SCRATCH to be controlled using pre-made template files in open-source drawing software (Inkscape already has plug-in support for many drawing-bots) (Fig S2; see also our shared template files). The user then loads an appropriate template for a given culture vessel, draws their desired patterns in each well, and ‘prints’ the scratch pattern on SCRATCH via a USB connection.

We demonstrated the versatility of SCRATCH by creating unique patterns in different types of Petri dishes and culture plates. First, we scratched a large-scale ‘star’ pattern across a layer of primary mouse skin keratinocytes in a 35 mm dish (Figure 1D) to demonstrate the ability to generate complex, precise patterns (see Fig. 1D, right). We then tested SCRATCH on a more challenging culture vessel – a 96-well plate. Here, the small well diameter prevents reproducible or precise manual scratching, and the throughput required to scratch 96-wells is not feasible using the traditional manual approach. However, SCRATCH was able to reliably pattern features (we used a ‘+’ shape) in all 96 wells in <4 minutes. Figure 1F shows a fluorescence image of the resulting patterns. Once calibrated, SCRATCH can automatically and reproducibly scratch arbitrary patterns in most standard culture dishes or plates at high throughput.

### Reproducibility and dynamics characterization

We first assessed how reproducible SCRATCH patterns were relative to manual patterns using linear scratches made in primary mouse skin keratinocyte layers cultured in 60 mm plates (see Methods); representative results are shown in Fig. 2A. We used the standard deviation of the width of each scratch as the metric for evaluating uniformity. As shown in Fig. 2B, SCRATCH exhibited significantly improved uniformity vs. manual scratching (nearly 4X reduction in standard deviation and on the order of a single cell), while maintaining an average width of ∼700 μm (approximate diameter of the 10 μL pipette tip). The observed variations we do see with SCRATCH likely reflect both biological variability in cell orientations and minor vibrations from the motor-belt system (see Fig S3 for high-resolution data on the tip trajectory, and Fig. S4 for a demonstration of the effective resolution limit).

**Figure 2.**
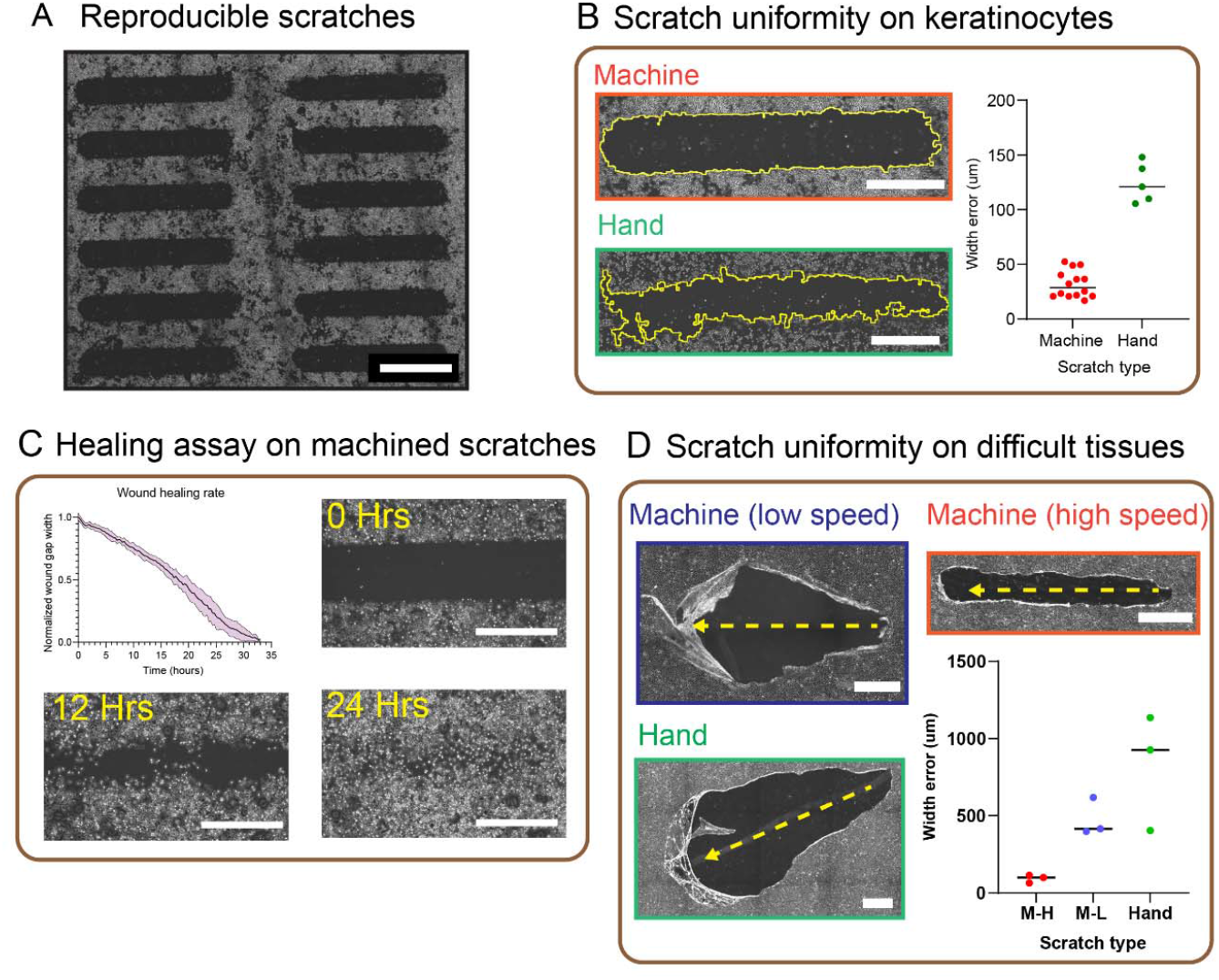
Linear scratch quantification and comparison. (A) 12 scratches performed by SCRATCH Scale bar: 2mm. (B) Scratch uniformity on keratinocyte monolayer. Edge outline is highlighted in yellow. Device scratch demonstrates lower with variation than manual. Scale bar: 1mm. (C) Wound healing assay on 8 scratches, showing uniform wound closure. Timelapse photos of 0, 12 and 24 hours after scratch are shown. Scale bar: 1mm. (D) Fast and consistent pipette movement from SCRATCH allows low scratch variation on high viscoelasticity tissues. A MDCK monolayer is scratched without calcium chelation. Scale bar: 1mm.

Therefore, SCRATCH demonstrates superior uniformity to manual scratches in basic tissues, which improves reproducibility of scratch assays and allows higher throughput. As a demonstration these benefits, we rapidly produced an array of 15 linear gaps into a primary mouse skin monolayer and quantified the wound closure rate to validate the uniformity (Fig. 2B). Phase-contrast images of 0 hour, 12 hours and 24 hours after scratching are shown alongside the quantification (Fig. 2C), and the closure curves indicate relative uniform and tight healing dynamics.

We next investigated the importance of scratching speed (how quickly the tool is translated through the tissue). This is something impossible to control manually, whereas SCRATCH allows scratch speed to be programmed up to 380 mm/s. Tissues are viscoelastic materials, meaning that their mechanical properties, adhesion to the substrate, and mechanobiological responses depend on the rate at which they are mechanically deformed, not just how much they are deformed, so being able to regulate the scratching rate should provide unique advantages and a new dimension to consider. In particular, we hypothesized that the high-speed, precise motion of SCRATCH would be particularly useful when working with more challenging tissues possessing strong cell-cell adhesion and relatively weaker cell-substrate adhesion where slow or irregular manual scratching can cause the tissues to delaminate rather than ‘cut’ (17).

Here, we used the widespread MDCK kidney epithelial model, commonly used in all manner of collective migration experiments and screens and known to exhibit strong cell-cell adhesion and develop collective cell behaviors as a result(9,18–21). We first established a baseline by manually scratching engineered, mature MDCK layers (see Methods) as best we could (Fig. 2D), which resulted in massive, irregular gaps and widespread delamination due to inherent irregularities in the manual process. We observed similar results when set SCRATCH to a slow speed (38 mm/s) and repeated the experiment (Fig. 2D). By contrast, when we repeated the experiment with SCRATCH to the fastest translation speed (380mm/s), we were able to produce highly uniform and more regular scratch patterns in comparison to slower mechanical or manual scratching (Fig. 2D). Overall, SCRATCH was able to deliver more precision, reproducibility, and throughput than manual scratching.

### Subtractive tissue manufacturing: designing complex tissue patterns

Only laboratory wounds are perfect straight lines, and many studies have emphasized the importance of tissue and wound shape in governing cellular migration and growth (10,22–27). We explored this concept by adapting SCRATCH for subtractive manufacturing of living tissues—gradually removing existing regions of tissue to produce complex patterns (returning to the primary mouse skin monolayer model). SCRATCH enables this by ‘raster cutting’, where it can gradually move the pipette tip tool back and forth while ensuring an overlap in the pattern to fully clear a given region of cells (Figs 3A-B). Here, we chose an approximate overlap of 75%. ‘Positive’ or ‘negative’ patterns can be achieved by selectively scratching the “center” or “edge” of a monolayer, either leaving a solid tissue (‘positive’) or cleared region (‘negative’) (Fig 3C). This subtractive manufacturing method extends the application of SCRATCH beyond pure scratch assays to complex assays evaluating the role of wound size and shape, for example. Moreover, this process is also fully automated within the free software used to control SCRATCH allowing arbitrarily complex patterns as shown in Fig. 3D.

**Figure 3:**
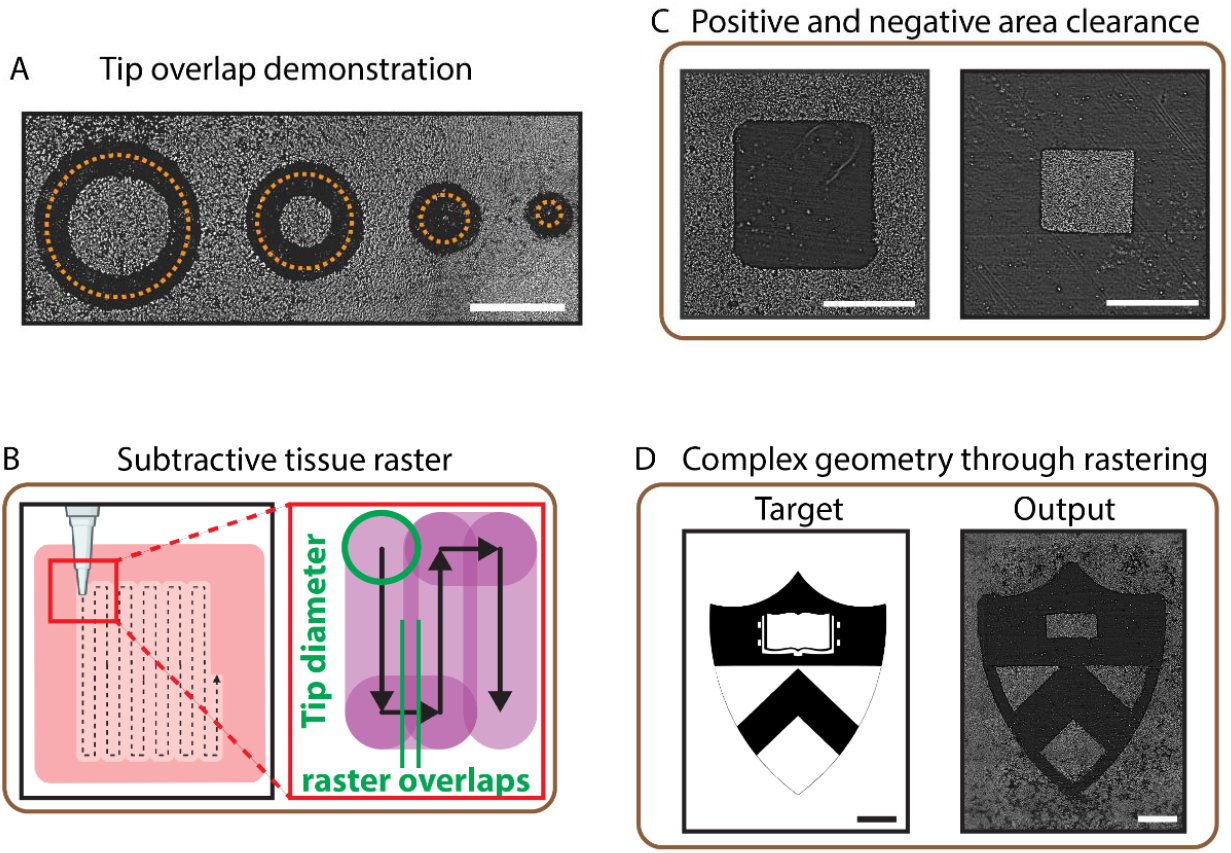
Raster mode capabilities and demonstration. (A) Demonstration of area clearance from tip overlap. The programmed path diameters are 3mm, 2mm, 1mm and 0.5mm. Scale bar: 2mm. (B) Raster mechanism cartoon and calculation of raster overlap. (C) Demonstration of positive and negative area clearance. Scale bar: 2mm. (D) Complex shape achieved through rastering. Scale bar: 2mm.

### SCRATCH for complex co-cultures

The “empty space” created by SCRATCH offers new potential for tissue co-culture because additional cell types can be back-filled into the newly created empty regions (Fig 4A). As a demonstration of this, we created a complex co-culture using a dermal/epidermal model of fibroblasts (3T3 fibroblasts) and keratinocytes (primary mouse keratinocytes). The resulting spiral pattern is shown in Fig. 4B-C and was produced by first scratching a layer of keratinocytes (pre-stained with a membrane dye), then washing with PBS and backfilling fibroblasts (pre-stained with a different membrane dye) as described in our Methods. The initial population of keratinocytes is shown in cyan and 3T3 in magenta. We also used a nuclear dye (Hoechst 33342) to stain all cell-types. The spiral is clearly visible and the expanded view shows good spatial separation between keratinocytes and fibroblasts. Note that the quality of the backfilling method relies on the confluency of the fist monolayer since the seeded cells will also attach to the area that is outside of intended region. Similar to planar lithography, this process can be repeated multiple times for additional “layers” of cells as long as a co-culture medium exists that can support each cell-type. These data further emphasize the versatility offered by the SCRATCH system to enable not only scratch assays but more complex tissue engineering and cell-cell communication assays.

**Figure 4.**
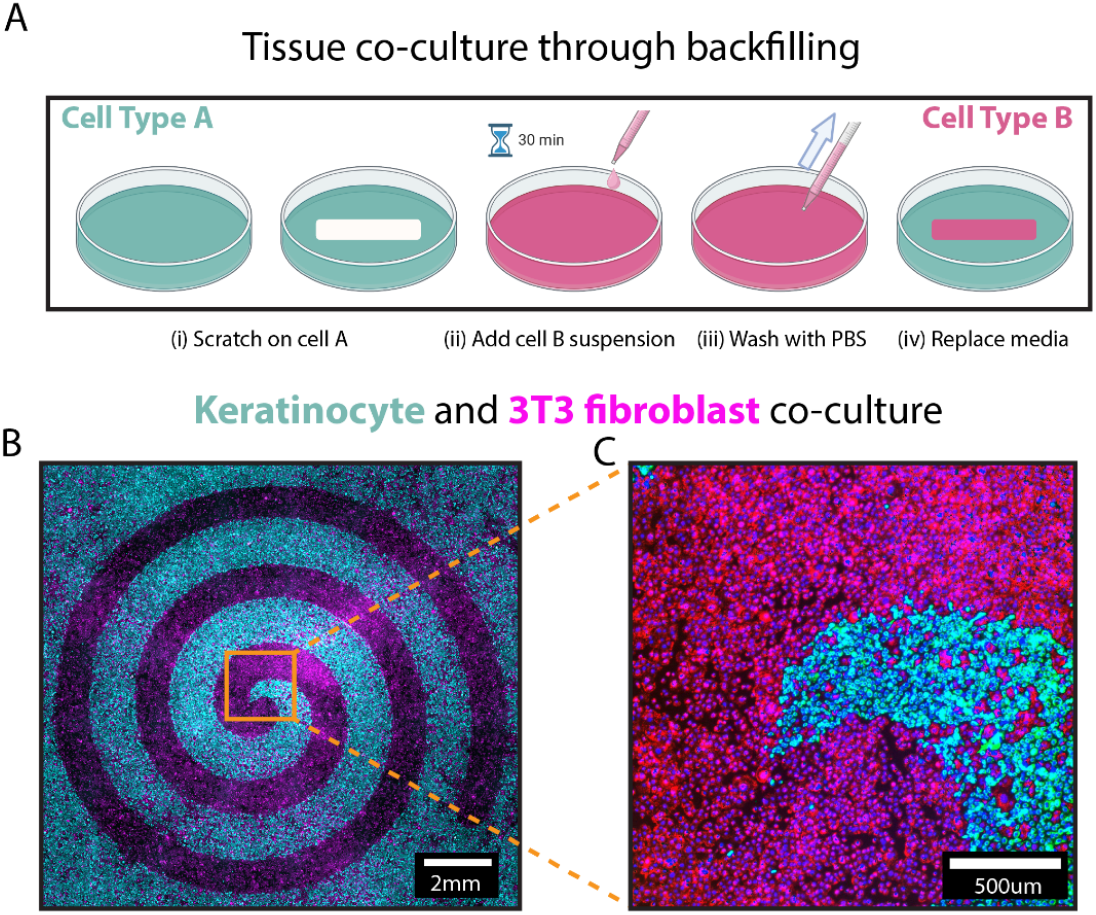
Using SCRATCH for arbitrary geometry tissue co-culture. (A) Tissue co-culture through backfilling. A scratch is made on Cell A monolayer. After washing with PBS, desired secondary cell suspension is added. After attachment, the dish is washed with PBS multiple times to remove any unattached cells. Co-culture media is then added and the dish is ready for experiment. (B) Fluorescence image of a spiral scratch on keratinocytes, then backfilled with 3T3 fibroblasts. Keratinocytes are stained with Cellbrite Green and 3T3 fibroblasts are stained with Cellbrite Red, both cells are stained with NucBlue. Scale bar: 2mm. (C) Zoomed in center of the spiral backfill. Scale bar: 500um.

## Discussion

SCRATCH demonstrates a low cost, fully programmable, and high throughput tool for the popular scratch assay that brings many significant advantages to the method, including improved reproducibility, throughput, and versatility with compatibility for nearly all standard culture plates and dishes. In particular, we showed improved precision, throughput, and reproducibility over manual scratches, as well as the ability to use scratching to produce unique tissue shapes and co-cultures without the need for microfabrication or manual stenciling.

The open-source and open hardware nature of SCRATCH, combined with its low cost, should substantially aid its adoption, as it can cheaply and easily be incorporated into most cell biological laboratories and used in or out of tissue culture hoods. A key aspect of SCRATCH is that it is easy to modify as a platform, allowing nearly any tip to be incorporated, and allowing for custom programming in Python if unique features are required that the standard graphics software does not allow (for instance, the tip can be programmed to go through a ‘wash’ step where it is agitated in a buffer or ethanol well in between scratching different wells in a multiwell plate to avoid cross-contamination). Similarly, the SCRATCH style platform can easily be modified with a more precise Z-drive to regulate scratching pressure, or enable tip-changes. Moreover, SCRATCH is not dependent on one specific piece of hardware, as any traditional ‘maker’ tool such as a diode laser cutter or 3D printer can be modified to do something similar. This type of versatility can substantially improve the types of applications where scratch-style assays are useful and further aid in its adoption.

## Methods

### Cell culture

Primary mouse keratinocytes were provided by the Devenport Laboratory at Princeteon University and cultured in E-medium (Nowak and Fuchs, 2009) supplemented with 15% serum (S11550, Atlanta Biologicals) and 50 μM calcium. Wild-type MDCK-II cells (courtesy of the Nelson Laboratory, Stanford University) were cultured in Dulbecco’s Modified Eagle’s Medium (D5523-10L, Sigma-Aldrich) with 1g/L sodium bicarbonate, 10% fetal bovine serum (S11550, Atlanta Biologicals), and 1% penicillin–streptomycin (15140-122, Gibco). NIH 3T3 fibroblasts were provided by the Schwarzbauer Laboratory at Princeton University. 3T3 cells were cultured in Dulbecco’s Modified Eagle’s Medium with phenol red (D5523-10L, Sigma-Aldrich), 10% fetal bovine serum (S11550, Atlanta Biologicals), and 1% streptomycin/penicillin (15140-122, Gibco). Tissue co-culture media consists of 50% Keratinocyte media and 50% 3T3 fibroblast media (28). All cells were maintained at 37 °C under 5% CO2 and 95% relative humidity. Cells were split before reaching 70% of confluence for maintenance culture, but all the dishes used for scratching had over 90% confluence to ensure even monolayers.

### SCRATCH hardware setup

Here, we used the Axidraw v3 drawing robot (Evil Mad Scientist, Inc.) to provide XYZ control of our scratching tip. All of the CAD files for the customized attachments and templates we describe here are available at our github repository (See Data Availability section). We designed and 3D printed a custom, modular plate holder that we attached to the Axidraw chassis using two M4 16mm long screws (94500A282, McMaster-Carr) and two M4 nuts (90592A090, McMaster-Carr), and this allows us to mount standard cultureware from 3.5 cm dishes to 96-well plates. We then designed and 3D printed a custom pipette holder with a thin layer of reusable adhesive (10079340647432, Loctite) (FIG S4). The pipette holder assembly was then gently clamped to the vertical stage of the Axidraw using the built-in clamping screw. We calibrated SCRATCH using an alignment ring around the target dish, and press-fit the dish into the modular plate holder. If needed, reusable adhesive can be added to improve stability. With the gantry in pen-up position and powered down (or its motors disengaged), we moved the gantry arm across the dish to ensure vertical clearance through the dish walls, and then aligned the pipette tip with the mark on the alignment ring, this establishes the “origin” of the drawing and the starting point.

Upon completion, the pipette tip holder assembly was removed from the vertical stage of Axidraw. Then the dish was removed from the holder and washed with PBS three times to remove cell debris.

### Scratch assay configuration

The Axidraw V3 is programmed using its official plugin in Inkscape (The Inkscape Team). The “Pen-up” and “Pen-down” range is set to 100% and 0% to ensure vertical clearance between the wells. Drawing speed is set to 10% (38mm/s) and pen-up movement speed is set to 75% (285mm/s). For contiguous tissues that have high cell-cell adhesions, drawing speed is set to 100% (380mm/s). Dialog box “Use constant speed when pen is down” is selected to ensure consistency. Pen raising speed and pen lowering speed is set to “Dead slow” to minimize pipette tip bouncing upon contact. Motor resolution is set to “∼2780DPI” for smooth operation and plot optimization set to least to avoid random starting point on a path. For all scratch assays, the programmed path is set to 0.01mm thick and is copied 4 times to the same place for repeated scratches. This ensures good area clearance and avoids uneven scratching due to non-conformal contact.

For raster mode, we use hatch fills options in Inkscape. Hatch spacing is set to a conservative value 0.1mm, which ensures each region is passed by the pipette tip at least 6 times to avoid any missed scratch zones due to non-conformal contact between the tip and the surface. Hatch angle is set to 45 degrees but can be modified based on the tip. Inset fill from edges option is selected to compensate for the finite tip width, and inset distance is set to 0.187mm (a 75% overlap to ensure path clearance) but should be determined experimentally.

### Tissue co-culturing

A 35mm dish with confluent keratinocytes was scratched with the steps shown previously. Then the dish was washed with PBS three times and stained with Cellbrite Green (30021, Biotium) at 5μL/mL for 30 minutes. A dish of 3T3 fibroblasts was also stained with Cellbrite Red (30023, Biotium) at 5μL/mL in suspension for 30 minutes. The stained dish is washed with PBS and 2ml co-culture media is added. Stained 3T3 suspension is washed with co-culture media 3 times using a centrifuge (5702, Eppendorf). 3T3 suspension is then added to the keratinocyte dish with a density of 1000 cells/mm^2. The dish is then incubated for 30 minutes for 3T3 attachment. Then the dish is fixed using 4% paraformaldehyde and stained with Hoechst 33342 (Thermo Fisher) for nucleus.

### Microscopy

Phase-contrast images were captured with an automated inverted microscope (Leica DMI8) with a 5X objective. For wound healing assays, time-lapse images were captured every 20 minutes. Fluorescence images were captured using an inverted microscope (Zeiss Axio Observer Z1) with a 5x objective, controlled using Slidebook (3I Intelligent Imaging Innovations) with Cy5, FITC, and DAPI filter sets. In both experiment setups, the microscopes were equipped with custom-built incubators maintaining 37 C and 5% CO2.

### Image and data analysis

FIJI (https://imagej.net/software/fiji) is used to process all images, including stitching (29) and wound area calculation for wound healing assay (MRI Wound Healing Tool, Montpellier Ressources Imagerie). Stitched phase images are processed through FFT bandpass filter (40px max, 2px min) to minimize flat fielding. A custom script is developed to analyze scratch width uniformity. Each scratch is thresholded and segmented to calculate the distance between the edges. Data visualization is performed using GraphPad Prism 10 (GraphPad Software).

## Data Availability

All CAD files for 3D printing and code necessary to perform the work shown here are available at our laboratory github repository (https://github.com/CohenLabPrinceton/SCRATCH) and we are happy to provide support as needed.

## Supporting Information

**Fig. S1: SCRATCH in operation**. Close-up image of SCRATCH operating on a 96-well plate. The tip is fixed using a thin layer of blue adhesive putty.

**Fig. S2: SCRATCH programming interface**. SCRATCH is programmed through Inkscape software. The dotted line represents scratchable area due to the contact angle between the tip and the edge of the dish. A star pattern is shown here in a 35mm dish template.

**Fig. S3: Path deviation of SCRATCH**. A pen filled with protein-A is fixed on the robot to show vibrations from the X- and Y-motors. The “wobble” deviation is 20um, significantly less than the pipet tip width of 700um.

**Fig. S4: SCRATCH resolution testing**. Scratch resolution using 10L tips.

Clearance is lost for scratches less than 1mm apart. Scale bar: 5mm

**Video S1: SCRATCH operation video**. A recording of SCRATCH in operation, accessing arbitrary wells and scratch a “cross” shape in a 96-well plate

## Acknowledgements

Support for this work was provided in part by NIH Award R35 GM133574-03. We also thank members of Cohen Lab for advice and support.

