## Supplemental information for "SCRATCH: A programmable, open-hardware, benchtop robot that automatically scratches cultured tissues to investigate cell migration, healing, and tissue sculpting"

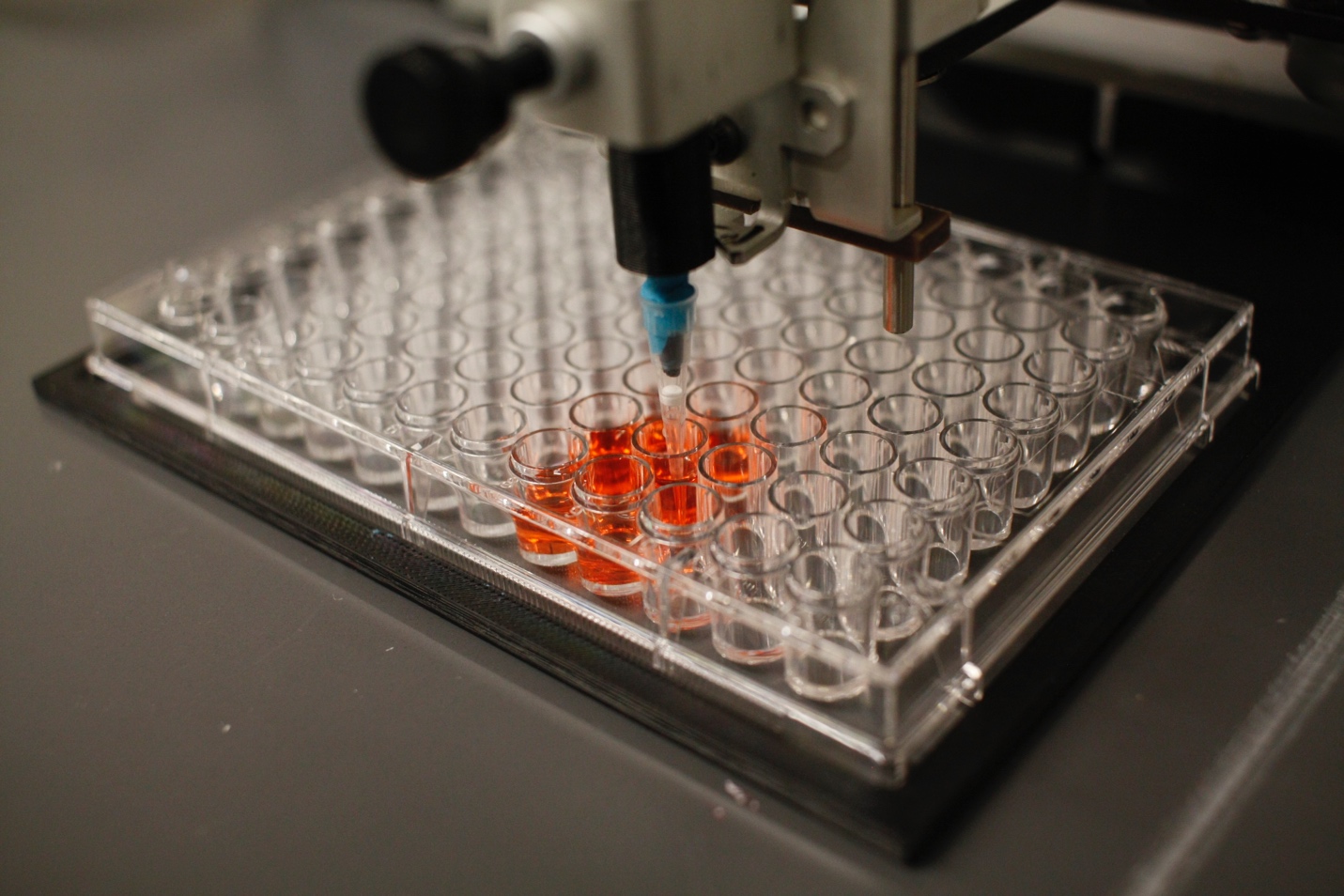

S1: Close-up image of SCRATCH operating on a 96-well plate. The tip is fixed using a thin layer of blue adhesive putty.

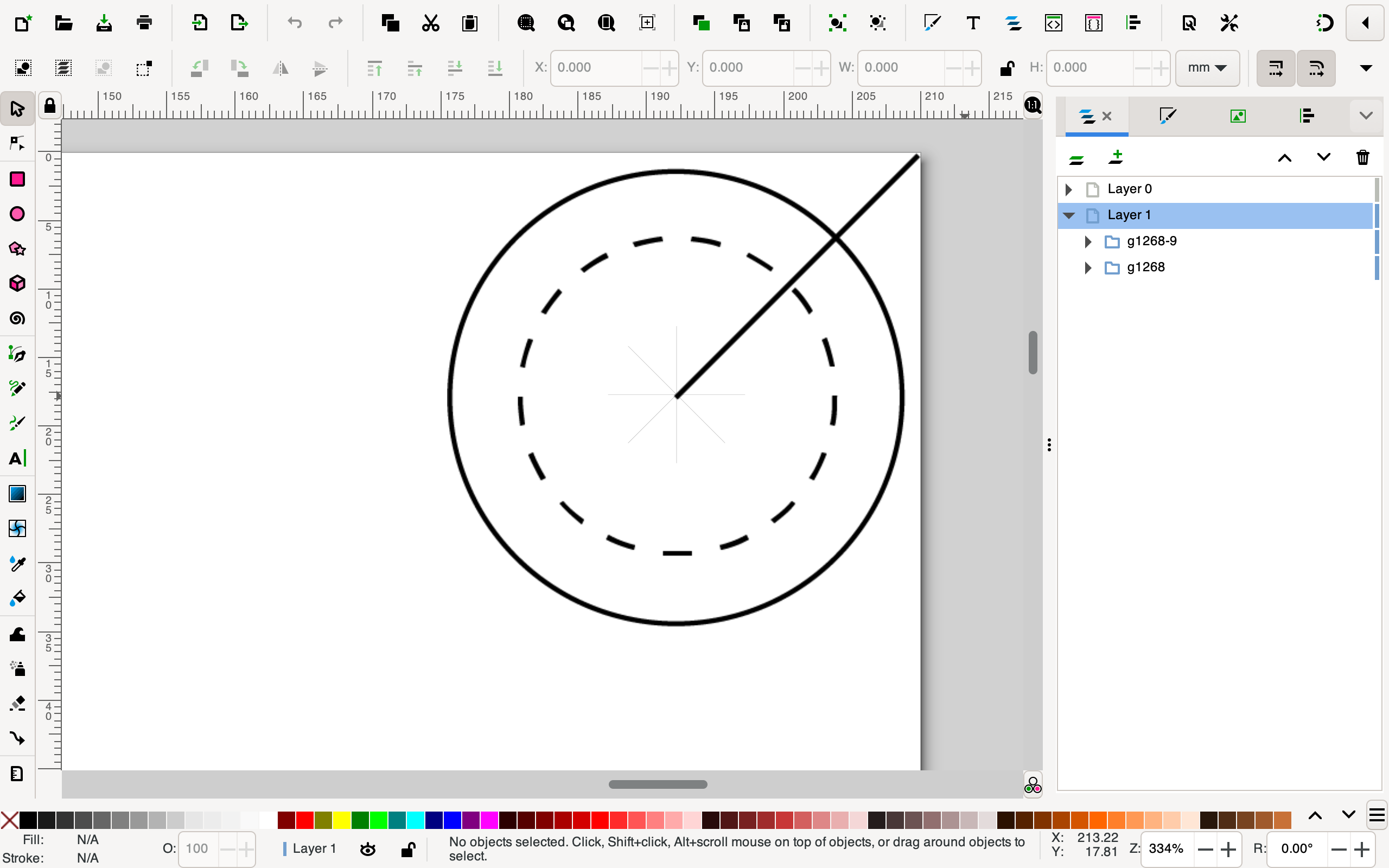

S2: SCRATCH programming interface through Inkscape software. The dotted line represents scratchable area due to the contact angle between the tip and the edge of the dish. A star pattern is shown here in a 35mm dish template.

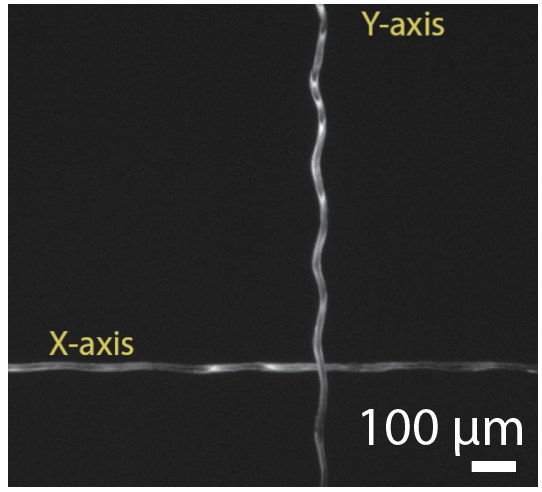

S3: A pen filled with protein-A is fixed on the robot to show vibrations from the X- and Y- motors. The “wobble” deviation is 20um, significantly less than the pipet tip width of 700um.

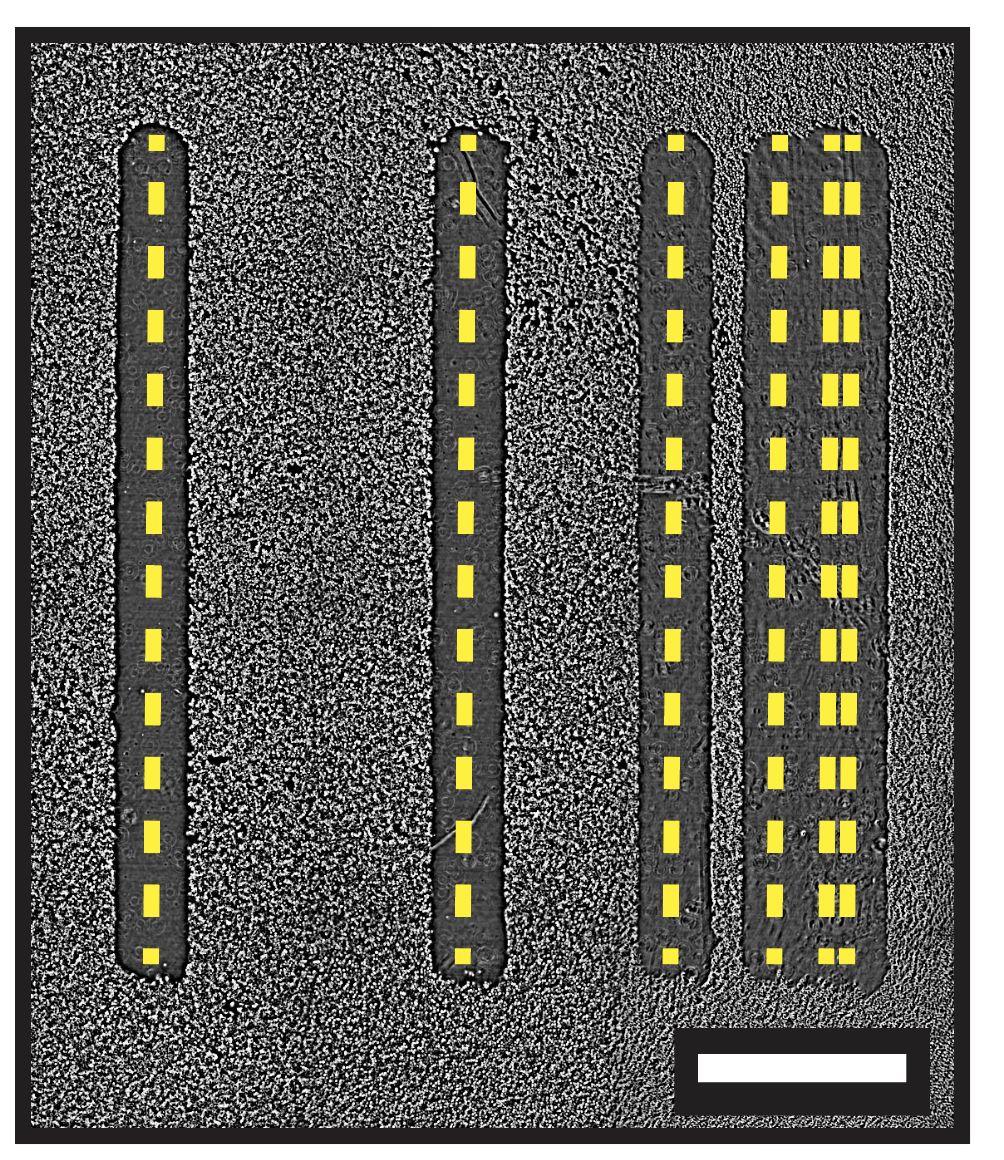

S4: Scratch resolution using 10L tips. Clearance is lost for scratches less than 1mm apart. Scale bar: 5mm
